# Sputum Production and Salivary Microbiome in COVID-19 Patients Reveals Oral-Lung Axis

**DOI:** 10.1101/2024.02.29.582705

**Authors:** Korina Yun-Fan Lu, Hend Alqaderi, Saadoun Bin Hasan, Hesham Alhazmi, Mohammad Alghounaim, Sriraman Devarajan, Marcelo Freire, Khaled Altabtbaei

## Abstract

**Objective:** SARS-CoV-2 is a severe respiratory disease that primarily targets the lungs and was the leading cause of death worldwide during the pandemic. Investigating the intricate interplay between the oral microbiome and inflammatory cytokines during the acute phase of infection is crucial for understanding host immune responses. This study aimed to explore the relationship between the oral microbiome and cytokines in COVID-19 patients, specifically examining those with and without sputum production.

**Methods:** Saliva and blood samples from 50 COVID-19 patients were subjected to 16S ribosomal RNA gene sequencing to analyze the oral microbiome. Additionally, 65 saliva and serum cytokines were assessed using Luminex multiplex analysis. The Mann-Whitney test compared cytokine levels between individuals with and without sputum production.

**Results:** Our study revealed significant differences in the membership (Jaccard dissimilarity: p=0.016) and abundance (PhILR dissimilarity: p=0.048; metagenomeSeq) of salivary microbial communities between COVID-19 patients with and without sputum production. Seven bacterial genera, including Prevotella, Streptococcus, Actinomyces, Atopobium, Filifactor, Leptotrichia, and Selenomonas, were present in statistically higher proportions of patients with sputum production (p<0.05, Fisher’s exact test). Eight bacterial genera, including Prevotella, Megasphaera, Stomatobaculum, Leptotrichia, Veillonella, Actinomyces, Atopobium, and Corynebacteria were significantly more abundant in the sputum-producing group, while Lachnoacaerobaculum was notably more prevalent in the non-sputum-producing group (p<0.05, ANCOM-BC).We observed a significant positive correlation between salivary IFN-gamma (Interferon-gamma) and Eotaxin2/CCL24 (chemokine ligand 24) with sputum production. Conversely, negative correlations were noted in serum MCP3/CCL7 (monocyte-chemotactic protein 3/Chemokine ligand 7), MIG/CXCL9 (Monokine induced by gamma/Chemokine ligand 9), IL1 beta (interleukin 1 beta), and SCF (stem cell factor) with sputum production (p<0.05, Mann-Whitney test).

**Conclusion:** Substantial distinctions in salivary microbial communities were evident between COVID-19 patients with and without sputum production, emphasizing the notable impact of sputum production on the oral microbiome and cytokine levels during the acute phase of infection.

## Introduction

COVID-19, an abbreviation for “coronavirus disease 2019,” originates from the SARS-CoV-2 virus and has exerted a global impact affecting millions [1]. While the initial wave of the pandemic may have receded, the persistence of new variants and the ensuing challenges underscores the imperative for sustained investigation into this novel pathogen. It is of vital importance to comprehend its multifaceted effects and develop innovative strategies to ameliorate the severity of clinical manifestations.

Host-specific characteristics exert a pivotal influence on the trajectory of SARS-CoV-2 infection, governing both its progression and severe outcomes [2]. The intricate interplay of local and systemic immune responses assumes a central role in shaping the host’s response to viral infection [2]. Predominant among the clinical expressions of COVID-19 are respiratory symptoms, encompassing cough, chest pain, dyspnea, and sputum production [1]. The viral route of entry, commencing through the oral cavity, followed by infection of respiratory epithelial cells and ensuing pulmonary inflammation, underscores the need to unravel the determinants governing these respiratory symptoms [3]. This necessitates a comprehensive exploration of local factors that explain immune-host responses such as the oral microbiome and inflammatory cytokines, which exert potential influence on symptom severity [3]. While existing literature highlights the connection between cytokine storms and severe manifestations [3], there exists a gap in terms of identifying specific microbiome and cytokine profiles linked to distinct COVID-19 symptoms. The present study endeavors to bridge this gap by directing its focus towards the oral microbiome and cytokine patterns relevant to sputum production—a significant clinical symptom in COVID-19.

Emerging research revealed a bidirectional interaction between COVID-19 and the oral microbiome [4]. Investigations have revealed that the oral immune system response may be influenced by the oral microbiome [4]. It was shown that patients infected with the virus exhibit distinctive oral microbial compositions compared to their non-infected counterparts [4]. This signifies the impact of the oral microbiota on the immune response during the acute phase of infection, thereby potentially contributing to the exacerbation of the clinical manifestation of COVID-19.

This study aimed to comprehensively elucidate the intricate interplay between the oral microbiome and cytokine profiles, especially those associated with respiratory symptoms, which is an area that remains incompletely explored. In this context, the present research aims to cover this gap. A primary objective involves the comparative analysis of oral microbiome compositions and cytokine profiles among individuals presenting with specific respiratory symptoms, with a particular emphasis on sputum production.

## Materials and Methods

This study was a collaborative joint study between the Harvard School of Dental Medicine (HSDM), J Craig Venter Institute (JCVI), the Ministry of Health in Kuwait, and the University of Alberta. The study was approved by JCVI, Harvard, the Kuwait Ministry of Health, and the University of Alberta. Informed consent was obtained from all enrolled participants (Kuwait Ministry of health ethics approval: #2020/1462 Harvard: IRB21-1492, University of Alberta: Pro00125245, JVCI: exempt due to secondary analysis of de-identified samples). All patients provided written consent to participation in the study.

### Study Design

A convenient sample strategy was used to recruit patients admitted at multiple Covid-19 centers in Kuwait between July 24th and September 4th, 2020. The data collection took place at three hospitals in Kuwait: Al Farwaniya Hospital, Jaber Al Ahmed Hospital, and Kuwait Field Hospital. Data were collected from those who provided positive consent and who tested positive for SARS-CoV-2 by RT-PCR from nasopharyngeal swabs (n=50). Saliva and serum samples were collected from individuals within 48 hours of PCR-confirmed COVID-19 diagnosis. The basic demographic and clinical information (including medical history, medication, weight, height, waist circumference, neck circumference, blood group, respiratory rate, and oxygen supplementation in liters for those on oxygen) of the study participants was obtained.

The severity of the disease was stratified into mild: hospitalized, no oxygen therapy (n=11); moderate: hospitalized, low-flow oxygen (<10 L/min) (n=28); and severe: hospitalized, high-flow oxygen (>10 L/min) (n=11). Mild and moderate groups were combined under “non-severe” in the present analysis.

### Saliva collection

15 mL plastic centrifuge tubes were prelabeled with the date and subject number. We then marked the 4 mL line of the tube. A parafilm was used to stimulate saliva.

Prior to sample collection, the saliva collection tube was placed in a cup with ice. A nurse, supervised by the research coordinator, would demonstrate how to provide saliva to the patient. Each subject was instructed to take a sip of water and rinse their mouth, swallow the water, chew the piece of parafilm, and then use their tongue to push saliva as it formed into the tube. They were then instructed to place the tube back in the cup with the ice cube while they waited for more saliva to form. The saliva was collected until it reached the line (4 mL) on the tube, considering the liquid region of the saliva sample (not the foamy regions). Once the patient finished providing the saliva sample, they notified the nurse. The nurse tightened the cap on the tube, wiped it with alcohol, placed the tube in the collection rack in the cooler with ice and discarded the other materials.

### Blood collection

Serum samples were collected using standard venipuncture techniques in 7.5 mL BD Vacutainer Serum marbled topped tubes with clot activator for serum. Samples were collected at the hospital for all samples.

### Sample processing

The samples were transferred to the Jaber Al Ahmad Hospital laboratory in containers with dry ice. The laboratory technician received the samples and processed them on the same day of sample collection within no more than 3 hours. Saliva samples were centrifuged at 2000 x g for 5 min, and the supernatants were separated from the pellets and transferred into different tubes. Serum samples were allowed to sit upright in racks at room temperature for 30 min prior to centrifuging at 2000 xg for 10 min at room temperature. All the samples were stored at –80 °C and were transferred from Kuwait to JCVI. The samples were placed on dry ice during shipment with a monitoring device to ensure that the samples were frozen during the transfer.

### DNA extraction, library preparation, and sequencing

DNA was extracted from samples using Qiagen’s AllPrep Bacterial DNA/RNA/Protein Kit (Cat# 47054; QIAGEN, Hilden, Germany) according to the manufacturer’s instructions. Step 3 was modified to use MP Biomedicals™ FastPrep-24™ Classic Bead Beating Grinder and Lysis System (Cat# MP116004500; FisherScientific) for 1 minute instead of vortexing for 10 minutes. All recommended products were used, and, at step 2, ß-ME was used instead of DTT.

Sequencing was done on the Illumina MiSeq platform. V4 hypervariable region was sequenced using the following forward (MSV4Read1) and reverse primers (MSV34Index1), 5’-TATGGTAATTGTGTGCCAGCMGCCGCGGTAA-3’ and 5’-ATTAGAWACCCBDGTAGTCCGGCTGACTGACT-’3.

Samples were deposited in NIH SRA under the accession number PRJNA948421.

### Cytokine abundance measurements

Serum samples were analyzed using the Immune Monitoring 65-Plex Human ProcartaPlex Panel (Cat# EPX650-16500-901; ThermoFisher Scientific, Vienna, Austria), according to manufacturer’s instructions, and runs on the Luminex 200 system (Luminex Corporation, Austin, Texas, USA). This kit measured immune function by analyzing 65 protein targets in a single well, including cytokine, chemokine, growth factors/regulators, and soluble receptors. The provided standard was diluted 4-fold to generate a standard curve, and high and low controls were also included.

### Amplicon sequence variants (ASVs)

Analysis was performed at the amplicon sequence variant level: DNA sequences containing no sequencing errors after algorithmic correction. The paired-end DADA2 [5] workflow of QIIME2 v2022.2.0 [6] was used for detecting ASVs and quantifying abundance implemented using the amplicon.py module of VEBA [7]. More specifically, this workflow uses the following protocol: 1) qiime tools import of paired-end reads; 2) DADA2 denoising of paired reads and ASV detection by qiime dada2 denoise-paired (forward_trim=251, reverse_trim=231, min-overlap=12); 3) taxonomic classification of ASVs using the precompiled silva-138-99-nb-classifier.qza model (Silva_v1383 SSURef_NR9); and 4) conversion into generalized machine-readable formats (e.g., QIIME2 Artifact and BIOM formatted files → tab-separated value and Fasta-formatted files).

### Statistical Analysis

Demographic data (Table 1) were collected, and descriptive statistics were computed for each variable, including mean and Standard Deviation (SD) for continuous variables, and counts and percentages for categorical variables. The chi-square test was utilized to assess the association between categorical variables, while the t-test was employed to evaluate differences in means between groups for continuous variables.

**Table 1.** Demographics and medical information of the study population.

| Variables |  | Sputum (n=23) |  | No sputum (n=27) |  | Total (n=50) | P value |
| --- | --- | --- | --- | --- | --- | --- | --- |
| Age, years, mean (SD) |  | 47.3 (12.0) |  | 53.8 (14.5) |  | 50.8 (13.7) | 0.1 |
| Sex | Female | 10 | 43.5 % | 12 | 44.4 % | 22 (44%) | 0.9 |
|  | Male | 13 | 56.5 % | 15 | 55.6 % | 28 (56%) |  |
| COVID-19 severity | Non-severe | 19 | 82.6 % | 20 | 74.1 % | 39 (78%) | 0.5 |
|  | Severe | 4 | 17.4 % | 7 | 26.9 % | 11 (22%) |  |
| Diabetes | Yes | 11 | 47.8 % | 11 | 40.7 % | 22 (44%) | 0.6 |
|  | No | 12 | 52.2<br>% | 16 | 59.3<br>% | 28<br>(56%) |  |
| **Heart**<br>**disease** | Yes | 2 | 8.7% | 6 | 22.2<br>% | 8<br>(16%) | 0.2 |
|  | No | 21 | 91.3<br>% | 21 | 77.8<br>% | 42<br>(84%) |  |
| **SOB** | Yes | 19 | 82.6<br>% | 13 | 48.1<br>% | 32<br>(64%) | **0.01\*** |
|  | No | 4 | 17.4<br>% | 14 | 51.9<br>% | 18<br>(36%) |  |
| **Smoking** | Yes | 1 | 4.3% | 0 | 0% | 1 (2%) | 0.3 |
|  | No | 22 | 95.7<br>% | 27 | 100% | 49<br>(98%) |  |
| **Chest**<br>**Pain** | Yes | 7 | 30.4<br>% | 5 | 18.5<br>% | 12<br>(24%) | 0.3 |
|  | No | 16 | 69.6<br>% | 22 | 81.5<br>% | 38<br>(76%) |  |
| **Asthma** | Yes | 4 | 17.4<br>% | 1 | 3.7% | 5<br>(10%) | 0.1 |
|  | No | 19 | 82.6<br>% | 26 | 96.3<br>% | 45<br>(90%) |  |
| **Emphyse**<br>**ma** | Yes | 0 | 0% | 1 | 3.7% | 1 (2%) | 0.4 |
|  | No | 23 | 100<br>% | 26 | 96.3<br>% | 49<br>(98%) |  |
| **WBC** |  | 5.8<br>(1.5) |  | 6.8<br>(3.7) |  | 6.3<br>(2.9) | 0.3 |

Sixty-five blood and salivary biomarker data (Table 2) were analyzed. For each biomarker, the median and interquartile range (IQR) were calculated for individuals with no sputum production (n=27) and those with sputum production (n=23). The non-parametric Mann-Whitney U test was utilized to assess the significance of differences between the two groups for each biomarker. A p-value threshold of less than 0.05 was considered statistically significant.

**Table 2.** Blood biomarkers and salivary biomarkers that are significantly different between individuals with and without sputum production.

|  |  | No sputum<br>(n=27), median<br>(IQR) | Sputum<br>(n=23),<br>median (IQR) | P<br>value |
| --- | --- | --- | --- | --- |
| **Blood**<br>**Biomarkers** | Eotaxin2/CCL24 | 711<br>(282, 1603.5) | 1325<br>(601, 1538) | 0.047 |
|  | MCP3/CCL7 | 11 (6,19) | 8 (6,11) | 0.042 |
|  | MIG/CXCL9 | 28.5 (1, 92) | 8.5 (-5, 43) | 0.011 |
|  | IL1 beta | -4 (-6,1) | -6 (-7,-5) | 0.011 |
|  | SCF | 34 (14, 131) | 14 (5,29) | 0.047 |
| **Salivary**<br>**Biomarkers** | IFNgamma | 0.688 (0.293,<br>1.111) | 1.245 (0.234,<br>2.813) | 0.047 |

IQR: interquartile range.

The aforementioned analyses were conducted using STATA SE 17.0 (StataCorp, College Station, TX, USA), with a significance level of p < 0.05 established for determining statistical significance.

### Microbiome statistical analysis

Bacterial composition data (Table 3) were analyzed using and subsequent data manipulation and adjustments were carried out using Microsoft Excel. To investigate significant differences in bacterial membership between the Sputum group (n=23) and the Control group (n=27), the following approach was taken. For each bacterial species, the presence of bacteria was represented as either a “1” (present) or a “0” (absent) for each subject. The number of “0s” (no bacteria) was then counted for each bacterial species in both groups. To assess the significance of differences in bacterial membership, a chi-square test was conducted for each bacterial species. The chi-square test determined whether the distribution of “0s” (absence of bacteria) significantly differed between the two groups. After obtaining chi-square test results from STATA, the subsequent data processing, adjustment of p-values using methods such as the Benjamin-Hochberg procedure, and table formatting were performed using Microsoft Excel. Adjusted p-values were calculated in Excel to control for multiple testing, and the final table was generated to present the adjusted p-values along with additional relevant information, including the number of subjects in each group with the bacteria (Table 3). Compositionally-aware differential abundance analysis was carried out using the ANCOM-BC [8] implementation in DATest package [9]. Beta diversity using PhILR [9] and beta-diversity was done on the automated pipeline FALAPhyl (available http://github.com/khalidtab/falaphyl).

**Table 3.** Bacteria that are significantly different in their membership between Sputum group and Control group. All are present with a significantly higher proportion in patients with sputum production.

| **Bacteria genera** | **Adjusted p value**<br>**(Benjamin-Hochberg)** | **number of subjects**<br>**with this bacterium**<br>**(Control group;**<br>**n=27)** | **number of subjects**<br>**with this bacterium**<br>**(Sputum group;**<br>**n=23)** |
| --- | --- | --- | --- |
| Prevotella | 0.04 | 1 (3.7%) | 5 (21.7%) |
| Filifactor | 0.03 | 1 (3.7%) | 9 (39.1%) |
| Actinomyces | 0.03 | 8 (3.0%) | 13 (56.5%) |
| Atopobium | 0.03 | 19 (70.4%) | 21 (91.3%) |
| Leptotrichia | 0.04 | 12 (44.4%) | 18 (78.3%) |
| Selenomonas | 0.04 | 12 (44.4%) | 18 (78.3%) |
| Streptococcus | 0.04 | 6 (22.2%) | 12 (52.2%) |

## Results

### Demographics

A total of 50 individuals who tested positive for SARS-CoV-2 were included in the study, consisting of 23 individuals with sputum production and 27 individuals without sputum production (Table 1). The average age for individuals with sputum production was 47.3, which was comparable to the average age of individuals without sputum production. The severity of the disease was categorized as mild (hospitalized without oxygen therapy, n=11), moderate (hospitalized with low-flow oxygen <10 L/min, n=28), and severe (hospitalized with high-flow oxygen >10 L/min, n=11). The analysis combined the mild and moderate groups into a “non-severe” category: 82.6% of the participants with sputum production exhibited non-severe symptoms and 17.4% exhibited severe symptoms. Among the subjects exhibited sputum production, 56.5% were males, and 43.5% were females. Diabetes was present in 47.8 % of the sputum positive group and 40.7% in the sputum-negative group. Heart disease was observed in 8.7% of the sputum-positive group and 22.2% of the sputum-negative group. 82.6% of those with sputum production experienced shortness of breath, which was significantly higher than the group without sputum production (48.1%). Chest pain was higher in those with sputum production, with 30.4% in the sputum-positive group and 18.5% in the sputum-negative group, but not to a statistically significant level. Asthma and emphysema were comparable between individuals with and without sputum production.

### Oral Microbiome

Our study showed significant differences in the membership (Jaccard dissimilarity: p=0.016) and abundance (PhILR dissimilarity: p=0.048; metagenomeSeq) of salivary microbial communities between COVID-19 patients with and without sputum production. In terms of membership, We found that seven bacterial genera were present in statistically higher proportions of patients with sputum production (p<0.05, Fisher’s exact test). These bacteria were: Prevotella, Streptococcus, Actinomyces, Atopobium, Filifactor, Leptotrichia, and Selenomonas. For abundance, eight bacterial genera were presented in significantly higher abundance in the group with sputum production, including Prevotella, Megasphaera, Stomatobaculum, Leptotrichia, Veillonella, Actinomyces, Atopobium, and Corynebacteria, and one genus, Lachnoacaerobaculum, was significantly more abundant in the group without sputum production (p<0.05, ANCOM-BC).

### Cytokines

We profiled saliva and serum samples to characterize oral mucosal and systemic responses following SARS-CoV-2 infection more comprehensively. Out of the 65 serum and salivary cytokines, five serum and one salivary cytokine were significantly different between individuals with and without sputum production. The serum Eotaxin2/CCL24 (chemokine ligand 24) and salivary IFN gamma (Interferon-gamma) were significantly elevated in individuals who reported sputum production, and MCP3/CCL7 (monocyte-chemotactic protein 3/Chemokine ligand 7), MIG/CXCL9 (Monokine induced by gamma/Chemokine ligand 9), IL1 beta (interleukin 1 beta), and SCF (stem cell factor) were found to be significantly lower in individuals who reported sputum production during the acute phase of COVID-19 infection (Table 2).

**Fig 1.**
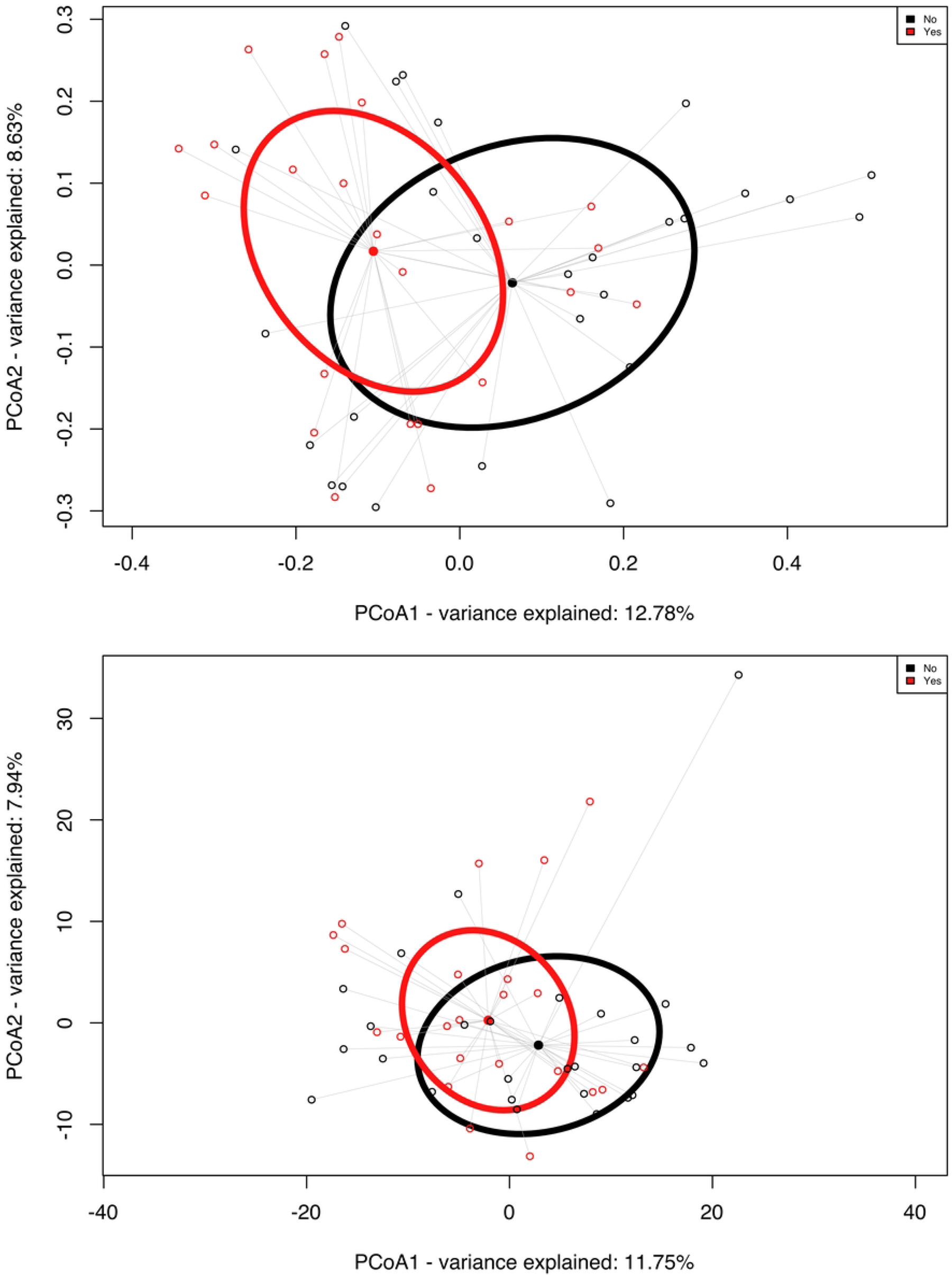
Analysis of salivary microbiome between the 2 groups, sputum positive (red, n=23) and sputum-negative (black, n=27). A:Principal coordinates analysis (PCoA) of Jaccard dissimilarity matrices between COVID-19 positive patients with sputum production and without sputum production. Ellipsoids show the deviation of spread of each group (ADONIS, p=0.016). B: PCoA of Euclidean distances of PhILR distances, demonstrating the three clusters between the conditions (ADONIS, p=0.048).

**Fig 2.**
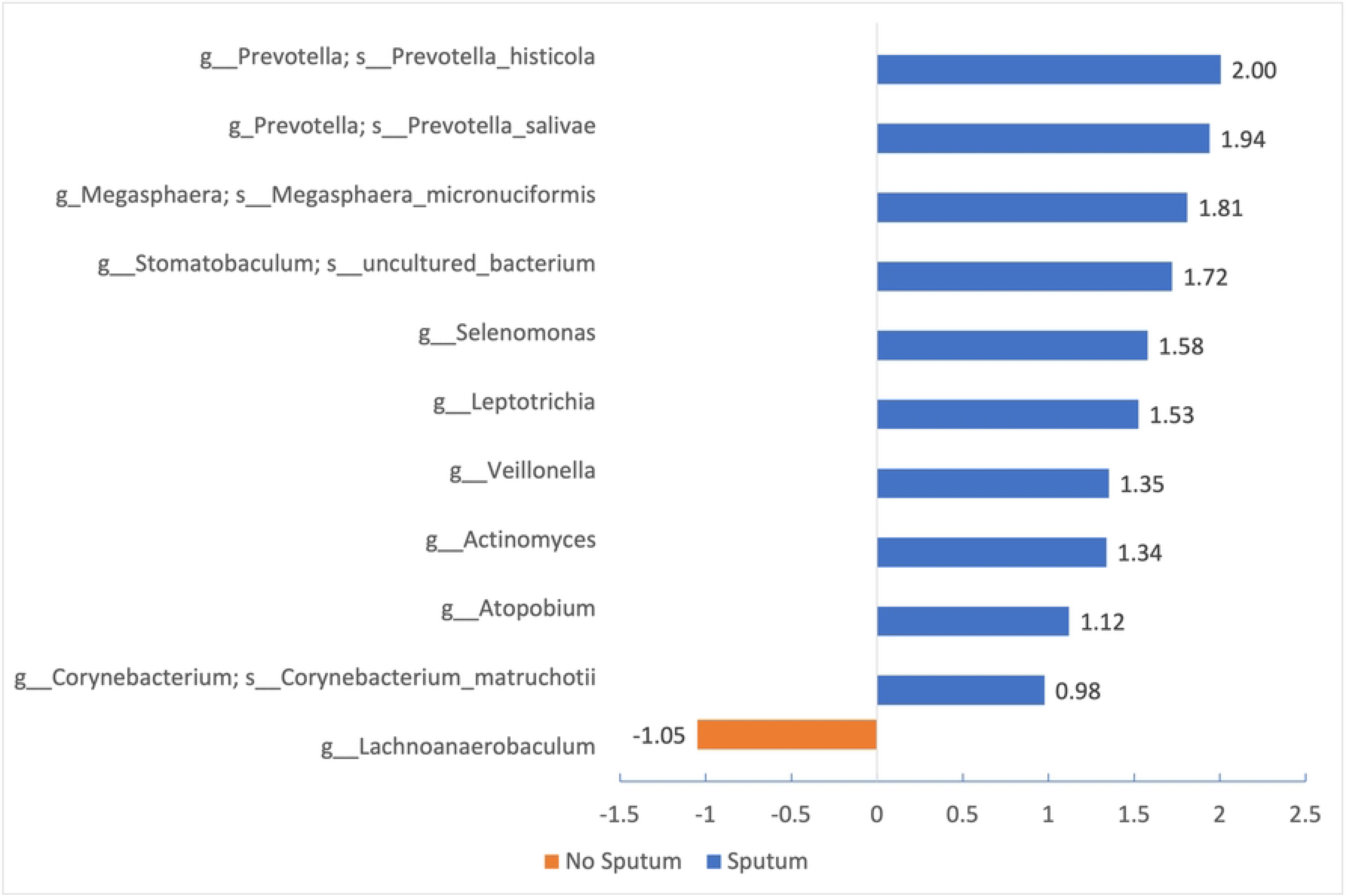
Differentially abundant bacteria (P<0.05, ANCOM-BC) in participants with Sputum production (blue) and without sputum production (orange).

## Discussion

### Oral microbiome

This study demonstrated a distinct bacterial cluster between people who developed sputum production compared to those who did not exhibit sputum symptoms during the acute phase of SARS-Cov-2 infection. Results showed distinct oral microbiome membership and abundance between patients with sputum production and controls. Seven bacteria genera were present in higher proportions of patients who exhibited sputum production: Prevotella, Filifactor, Actinomyces, Atopobium, Leptotrichia, Selenomonas, and Streptococcus. Additionally, there was a major difference in the abundance of bacteria between those who manifested sputum production compared to those who didn’t. Eight bacterial genera exhibited significantly higher levels in the group with sputum production. These included Prevotella, Megasphaera, Stomatobaculum, Leptotrichia, Veillonella, Actinomyces, Atopobium, and Corynebacteria. Conversely, one genus, Lachnoacaerobaculum, was notably more abundant in the group without sputum production. Notably, Prevotella, Actinomyces, Atopobium, and Leptotrichia displayed differences both in their membership and abundance.

The identification of distinct bacterial clusters and differences in oral microbiome membership and abundance between COVID-19 patients with and without sputum production has significant implications, particularly when considering the potential connections between these findings and oral health conditions, particularly periodontitis. Prevotella, Atopobium, Filifactor, and Actinomyces, for instance, have well-established connections with periodontal health and oral infections [10-12]; Leptotrichia species can exhibit invasive traits, potentially causing various infections, and Megasphaera species are linked to periodontal disease and polymicrobial infections [13,14]. The association of these bacteria to periodontal diseases is of significance as periodontitis has been shown to be associated with an increased likelihood of developing COVID-19 symptoms [15,16]. Individuals with underlying periodontal issues might face an increased susceptibility to COVID-19 complications due to oral-systemic links [16]. Emerging research has revealed a potential association between moderate-to-severe periodontitis and heightened risks of COVID-19 complications, including severe outcomes such as death, ICU admission, and the need for assisted ventilation [15]. These findings suggest that periodontal disease may exacerbate the severity of COVID-19 infections, possibly due to shared inflammatory pathways and risk factors with chronic inflammatory conditions known to influence COVID-19 outcomes [15]. Although we didn’t perform oral exams on the individuals in our study, it is essential to consider the potential impact of periodontal health on respiratory symptoms like sputum production, as it has been associated with an increased likelihood of COVID-19 symptoms [15].

We propose two potential pathways that periodontitis could be linked to COVID-19 respiratory symptoms such as sputum production: periodontitis is recognized as a chronic inflammatory condition, and this inflammation could potentially exacerbate the severity of COVID-19 symptoms [15]. Secondly, periodontal pathogens, including specific bacteria like Prevotella, Atopobium, Filifactor, Actinomyces, Leptotrichia, and Streptococci, may have the capacity to migrate into the respiratory tract during a COVID-19 infection, causing respiratory symptoms like sputum production [10,11,13,17].

The temporal origin of these bacteria identified in sputum, whether existing prior to COVID-19 infection and exacerbating symptoms such as sputum production, or emerging during the infection’s progression, remains unclear. Previous studies documented that in the sputum samples of healthy subjects, Firmicutes, Bacteroidetes and Actinobacteria were the major phyla, and Streptococcus, Veillonella, Prevotella, Haemophilus, Actinomyces and Rothia were among the dominant genera [18,19]. Therefore, a hypothesis can be formulated, suggesting that the oral disease-related bacterial species we identified in individuals with sputum production might have already been present prior to COVID-19 infections due to specific periodontal diseases or dental caries. These bacterial species could potentially contribute to exacerbating respiratory symptoms and intensifying inflammation. This underscores the significant role of oral health as a potential gateway to systemic diseases. Notably, a previous analysis of the same dataset unveiled marked differences in oral microbiome profiles between individuals with severe symptoms and those with mild symptoms [20]. This suggests the potential impact of the oral immune system and microbiome on the severity of COVID-19 symptoms.

### Cytokines

Our analysis demonstrated that certain cytokines were associated with sputum production. Out of the 65 serum and salivary cytokines we analyzed, serum Eotaxin2/CCL24, and salivary IFN-gamma were elevated in individuals who reported sputum production, and MCP3/CCL7, MIG/CXCL9, IL1 beta, and SCF were found to be lower in individuals who reported sputum production during the acute phase of COVID-19 infection.

These findings are particularly intriguing when considering the role of cytokines in immune responses in the context of COVID-19 and oral dysbiosis. Strong evidence showed that the oral microbiome plays a crucial role in the development and maintenance of the immune system’s homeostasis [21]. Oral dysbiosis, characterized by an imbalance or perturbation in the composition of the oral microbiome, profoundly influences the host immune response and the intricate network of cytokines involved [22]. When the oral microbiome experiences dysregulation, often associated with conditions like periodontal disease, the immune system mounts a multifaceted response [23]. This immune reaction involves the secretion of various cytokines, which are typically upregulated in response to oral dysbiosis, driving inflammation to combat the overgrowth of pathogenic bacteria [23]. Many proinflammatory cytokines, including IFN-gamma [24,25] and Eotaxin2/CCL24 [26], have been shown to be correlated with worse clinical outcomes associated with COVID-19. The heightened levels of these two cytokines in individuals with sputum production may suggest an overly robust immune response, potentially exacerbating disease severity. As we previously proposed, the respiratory symptoms of COVID-19 could be intercorrelated with the complex interplay between periodontal diseases, oral dysbiosis, and cytokine dysregulation. Moreover, cytokines MCP3/CCL7, MIG/CXCL9, IL1 beta, and SCF, key in viral infection immune responses, exhibited reduced levels in COVID-19 patients with sputum production during the acute phase. This suggests potentially less effective immune defense against the virus. MCP3/CCL7 draws monocytes, T cells, and dendritic cells to infection sites [27], while MIG/CXCL9 attracts T cells and natural killer cells [28]. Pro-inflammatory cytokine IL-1 beta initiates immune responses [29], and SCF supports white blood cell, including T cell, production [30]. Lower cytokine levels might imply a compromised immune reaction, potentially resulting in severe infections or extended recovery.

## Limitations

This study employed a cross-sectional design, and the relatively modest sample size could potentially limit our capacity to detect significant differences among different presentations of respiratory symptoms. Additionally, due to pandemic-related circumstances, we were unable to gather data on periodontal health and dental caries, despite the potential influence of oral health status on the salivary microbiome. It’s important to acknowledge the limitations inherent in our study, emphasizing the necessity for subsequent research endeavors. We also acknowledge that any definitive connection between these taxa and sputum production should be verified with actual sputum samples from within the respiratory cavity.

## Conclusion

There was notable distinction in the oral bacterial composition and abundance, alongside levels of inflammatory cytokines in both serum and saliva, between patients exhibiting sputum production and those without it. These variations suggest distinct host immune responses between the two groups. Particularly noteworthy is the heightened presence and abundance of numerous pathogenic bacteria associated with oral diseases in individuals experiencing sputum production. Considering the oral cavity as the initial point of entry for COVID-19, an intriguing hypothesis takes shape: a compromised oral health could potentially act as a contributing factor, enhancing the likelihood of respiratory symptoms linked to SARS-CoV-2, notably including the production of sputum.

## Acknowledgements

We acknowledge the funding agencies for the support to this work.

